# PAIRUP-MS: Pathway Analysis and Imputation to Relate Unknowns in Profiles from Mass Spectrometry-based metabolite data

**DOI:** 10.1101/209577

**Authors:** Yu-Han H. Hsu, Claire Churchhouse, Tune H. Pers, Josep M. Mercader, Andres Metspalu, Krista Fischer, Kristen Fortney, Eric K. Morgen, Clicerio Gonzalez, Maria E. Gonzalez, Tonu Esko, Joel N. Hirschhorn

**Author notes:** Corresponding author: Joel Hirschhorn.

## Abstract

Metabolomics is a powerful approach for discovering biomarkers and metabolic quantitative trait loci. While untargeted profiling methods can measure up to thousands of metabolite signals in a single experiment, many signals cannot be readily identified as known metabolites or compared across datasets, making it difficult to infer biology and to conduct well-powered meta-analyses across studies. To deal with these challenges, we developed a suite of computational methods, PAIRUP-MS, to match metabolite signals across mass spectrometry-based profiling datasets using an imputation-based approach and to generate pathway annotations for these signals. We performed meta and pathway analyses for both known and unknown signals in multiple datasets and then validated the results using genetic associations. Finally, we applied the methods to detect metabolite signals and pathways associated with body mass index, demonstrating that our framework is useful for analyzing unknown signals in a robust and biologically meaningful manner and for improving the power of untargeted metabolomics studies.

## INTRODUCTION

Metabolomics is a powerful systematic approach for studying circulating molecules in biological samples^1,2^. Profiling technologies such as nuclear magnetic resonance (NMR) spectroscopy and liquid or gas chromatography followed by mass spectrometry (LC-MS or GC-MS) have been used to discover metabolites that are biomarkers or predictors for various human traits and diseases, including clinical risk factors of metabolic or cardiovascular diseases^3^, type 2 diabetes^4^, and all-cause mortality^5^. Genome-wide association studies (GWAS) have also identified numerous metabolic quantitative trait loci (mQTLs) that influence variation in metabolite abundance, which may provide valuable insights into gene functions and disease mechanisms^6–9^.

While metabolomics has led to many new discoveries, a significant portion of the generated data have not been used to its full potential. Untargeted profiling platforms collect measurements for thousands of metabolite signals and compare their data to standards in compound databases to identify known metabolites^1^. Current identification procedures can identify up to a few hundred known metabolites and the remaining unknown signals are often excluded from downstream analyses. However, studying unknown signals may be fruitful for several reasons. First, many unknowns show correlation with known metabolites, phenotypes, and genetic variants, indicating that they capture biological information^10^. Second, because unknowns usually make up a large portion of untargeted data, including them in analysis can greatly increase the search space in analyses. Finally, investigating unknown signals may enable us to discover novel biomarkers that have not been characterized in human metabolism. Therefore, developing computational methods that deal with unknown signals is critical for making the most out of untargeted metabolomics data.

Analyzing unknown signals poses several challenges. Distinguishing biological signals from noise and adjusting for technical artifacts are difficult, because unknowns are often associated with noisier measurements compared to known metabolites^11^. Meta-analysis, which may increase power for detecting noisy but true signals, is complicated by the lack of gold standard protocols and varying signal characteristics across platforms and studies. While some methods have been developed to align signals across datasets using mass-to-charge ratio (m/z) and/or retention time (RT) information, their applications are often restricted to processing data batches within a single study or data generated by the same profiling method^11–13^. Another main challenge is to link unknown signals to biological functions without confirming their chemical identities. Although several pathway or network-based approaches have incorporated unknown signals into metabolomics analyses, they focus on a small subset of the unknowns (e.g. those directly associated with known metabolites, genes, or a trait of interest) and do not explicitly provide biological interpretations (e.g. pathway labels) for enriched pathways or network clusters identified during analysis^10,14,15^.

To overcome the challenges described above, we developed and implemented computational methods for analyzing unknown signals. Our suite of methods, called PAIRUP-MS (Pathway Analysis and Imputation to Relate Unknowns in Profiles from MS-based metabolite data), can be used to process, match, and annotate metabolite signals in MS-based profiling datasets and to perform meta and pathway analyses for these signals. We utilized genetic associations to validate the methods and demonstrated their application by studying metabolite signals associated with body mass index (BMI). Our framework can be applied to diverse untargeted profiling datasets and will help to advance metabolomics as a powerful approach for elucidating biology underlying human traits and diseases.

## RESULTS

### Metabolomics datasets and data processing

We developed and tested computational methods in PAIRUP-MS using LC-MS profiling data generated from three independent cohorts: (1) Obesity Extremes cohort (OE): 300 individuals (100 lean, 100 obese, and 100 population control) sampled from the Estonian Biobank, (2) Mexico City Diabetes Study cohort (MCDS): 865 individuals in a prospective type 2 diabetes study, and (3) BioAge Labs Mortality cohort (BioAge): 583 individuals selected from the Estonian Biobank for a retrospective mortality study. In order to reduce noise in the profiling data and to improve power in analyses, we implemented a quality control (QC) pipeline, described in **Methods** and **Supplementary Information**, to adjust each dataset for measurement variation associated with technical artifacts and remove samples, metabolite signals, and data points with noisy trends. After QC, we leveraged the correlation structure among signals to impute any remaining missing values within a dataset. In the end, the OE dataset contained 298 samples and 13,613 metabolite signals (322 known and 13,291 unknown); MCDS contained 821 samples and 7,136 signals (242 known and 6,894 unknown); BioAge contained 583 samples and 14,617 signals (603 known and 14,014 unknown). Covariate adjustment (if appropriate) and rank-based inverse normal transformation were performed on each metabolite signal and the resulting abundance *z*-scores were used in downstream analyses. QC’ed cohort and dataset characteristics are summarized in **Table 1**.

**Table 1.** Characteristics of study cohorts and corresponding metabolomics datasets

|  | **Obesity Extremes cohort (OE)** | **Mexico City Diabetes Study cohort (MCDS)** | **BioAge Labs Mortality cohort (BioAge)** |
| --- | --- | --- | --- |
| **Study design** | BMI extremes from Estonian Biobank (EB) | Prospective cohort for type 2 diabetes | Retrospective cohort for mortality from EB |
| **# of samples** | 298 | 821 | 583 |
| **# of women (%)** | 149 (50.0) | 498 (60.7) | 407 (69.8) |
| **Age** | 38.5 ± 12.1 | 52.3 ± 7.73 | 73.3 ± 2.70 |
| **BMI** | 28.3 ± 9.95 | 28.6 ± 4.54 | 27.3 ± 4.33 |
| **Fasting time\*** | 7.42 ± 3.34 | overnight | 3.04 ± 2.80 |
| **Profiling methods\*\*** | *C8-pos, C18-neg, HILIC-pos, HILIC-neg* | *LIPID, HILIC-pos, CMH* | *C8-pos, C18-neg, HILIC-pos, HILIC-neg* |
| **# of signals (# of knowns)** | 13,613 (322) | 7,136 (242) | 14,617 (603) |
| **# of signals with m/z data (# of knowns)** | 13,552 (261) | 7,083 (189) | 14,617 (603) |

Mean ± standard deviation is reported for quantitative phenotypes. \* Fasting time for OE and BioAge was not exact due to conversion from categorical variable; 200 BioAge samples had no fasting time information. \*\* Italicized profiling methods had m/z and RT information available; we treated "C8-pos" and "LIPID" to be the same method in our analyses due to large overlap in the metabolites they measured; see **Supplementary Information** for methods details.

### Matching metabolite signals across datasets

#### Matching method overview

An approach for comparing unknown signals measured in different untargeted profiling datasets is crucial for increasing power in metabolomics studies. Instead of using RT information, which can vary greatly across profiling methods and platforms, we developed a method that utilizes m/z and imputation to match up signals likely to represent the same metabolites across MS-based profiling datasets (**Fig. 1a**). In preliminary analysis, we observed that known metabolites measured across our three datasets had moderately similar correlation structures despite substantial differences in study populations and designs (**Supplementary Fig. 1**). We took advantage of this observation and imputed the abundance of unknown (or unshared known) signals from one dataset to another using shared known metabolites as predictors in linear regression models. The imputation then allowed us to calculate the correlation between signals measured in different datasets across a common set of samples and to pair up signals based on agreement in m/z and this correlation.

**Figure 1.**
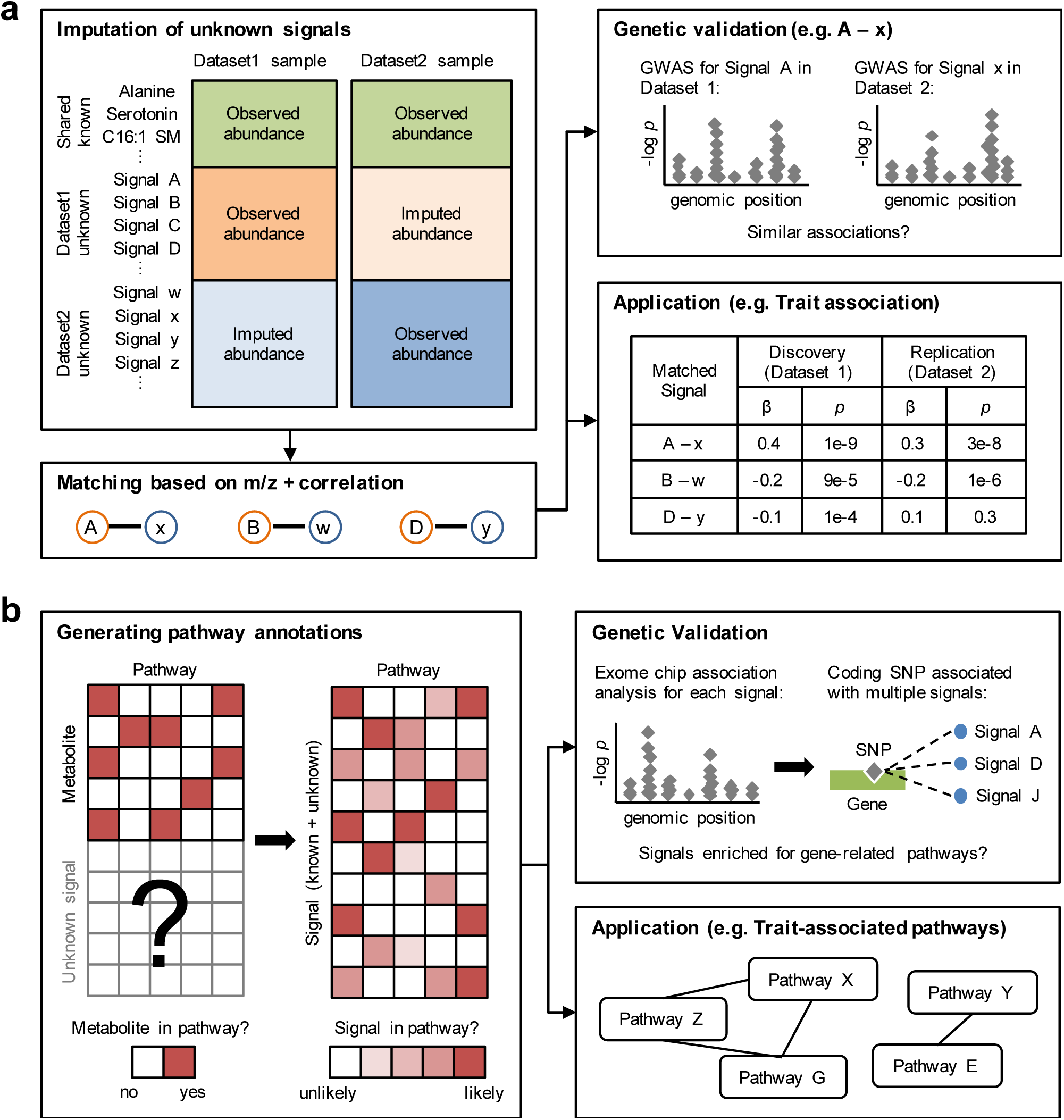
PAIRUP-MS is a suite of computational methods for analyzing metabolite signals in untargeted metabolomics data. (a) Overview of matching method: unknown (or unshared known) signals are imputed across datasets using shared known metabolites as predictors and then paired up based on m/z and correlation across samples. Genetic data can be used to validate matched signal pairs that share similar genetic associations. Matched signals can be used to perform combined association analyses across datasets (e.g. identifying trait-associated signals in discovery and replication cohorts). (b) Overview of pathway method: using binary metabolic pathway annotations and untargeted profiling data as input, a pathway reconstitution procedure is performed to construct a numeric annotation matrix, where each signal (known or unknown) gets a membership score in each pathway (or “metabolite set”, see Methods). Genetic data can be used to validate that signals associated with a specific gene are enriched for reconstituted pathways related to the gene. The annotation matrix can be used to perform pathway analyses (e.g. identifying pathways enriched for a list of trait-associated signals).

#### Calibration using shared known metabolites

In order to calibrate parameters in our method, we treated known metabolites shared across two datasets as unmatched signals and applied our method to match each of them in turn. We compared the performance of different parameter settings by calculating the percentage of matches that were either the correct match or highly correlated (r^2^ > 0.8) with the correct match in observed data. The parameter choices we tested included: (1) whether to consider different adduct ions when checking for m/z agreement, (2) which set of samples to use for calculating correlation between signals, (3) whether to allow multiple-to-1 signal matching or require unique 1-to-1 matching, (4) whether to match only signals measured by the same profiling method or allow matching across different methods, and, optionally, (5) which correlation cutoff threshold to use for calling a match.

The calibration results for OE-MCDS, OE-BioAge, and MCDS-BioAge matching are shown in **Supplementary Figs. 2-4** and the optimal parameter settings chosen for each dataset pair are summarized in **Supplementary Table 1a**. We decided to consider two types of matches, “multiple” and “reciprocal”, for subsequent analyses. Briefly, in the “multiple” setting, we matched each Dataset 1 signal, *s*, to a Dataset 2 signal by first identifying all Dataset 2 signals whose m/z agree with *s*, and then selecting the one that has the best correlation with *s* to be the final match. Following this procedure, more than one Dataset 1 signal may be mapped to the same Dataset 2 signal. For “reciprocal” matches, we repeated the same procedure to match Dataset 1 signals to Dataset 2 and vice versa, and only kept the subset of matches that were generated in both directions. Our method performed slightly better using the “reciprocal” setting, in which it correctly paired up 69.9 to 87.6% of the matched knowns, with 91.6 to 96.4% of all matches showing strong correlation (r^2^ > 0.8) with the correct match. However, the “multiple” setting also performed relatively well (58.4 to 87.1% of matches were correct; 91.0 to 96.1% were highly correlated) and identified more correct or highly correlated matches overall, so both match types can be useful, depending on the analysis context.

We compared our method against a m/z and RT-based matching approach, where signals with agreeing m/z were mapped to each other based on predicted RT calculated from shared known metabolite data. The imputation method generally performed as well as or slightly better than the RT method, despite not using any RT information (**Supplementary Table 1a**). An important advantage of the imputation approach is that, even when the matches made for known metabolites were incorrect, ~60-80% of them were strongly correlated (r^2^ > 0.8) with the true known matches. In contrast, incorrect matches made by the RT approach sometimes showed much weaker correlation. These results suggest that our method can be used to discover “proxies” for metabolites in a dataset even when an exact match does not exist or could not be identified. Furthermore, when we allowed matching across profiling methods, the RT approach showed a bigger dip in performance compared to the imputation approach (**Supplementary Table 1b**). Thus, imputation-based matching may be a more flexible framework for comparing datasets generated by diverse profiling platforms, especially when directly comparable RT data is not available.

#### Genetic validation of matched signals

After demonstrating that the matching method could be used to accurately pair up shared known metabolites, we next sought to validate the full set of matches. Because there is no uniform “gold standard” for matching unknown signals, we used genetic data to validate matched signals between OE and MCDS, reasoning that useful matches would have similar patterns of genetic association across the two cohorts. We performed a genome-wide association study (GWAS) in both OE and MCDS for the matched signal pairs (4,432 “multiple” matches, of which 1,573 of which were “reciprocal”). We also analyzed an additional 207 pairs of shared knowns and 20 sets of 4,432 randomly matched pairs to serve as positive and negative controls, respectively. To assess how well we had selected matching pairs, we determined the overall directional consistency of association for the best-associated single nucleotide polymorphisms (SNPs) with the matched signal pairs. Specifically, for each matched pair, we identified the SNP with the best *p*-value in a GWAS meta-analysis of OE and MCDS, and tested whether the same allele was associated with increased metabolite levels across the two studies. (To avoid biasing the subsequent assessment of directional consistency, we forced all SNPs to have the same direction of effect when calculating meta-analysis *p*-values; see **Methods**).

For the 94 pairs of shared knowns that had genome-wide significant (*p* < 5 × 10^−8^) SNPs in the meta-analysis, 66 pairs (70.2%; Fisher’s *p* = 1.56 × 10^−4^) showed directional consistency; in comparison, 478 out of 672 (71.1%; Fisher’s *p* = 1.83 × 10^−28^) “reciprocal” or 1,119 out of 1,772 (63.1%; Fisher’s *p* = 3.08 × 10^−28^) “multiple” matched pairs were consistent, both of which were significantly better than randomly matched signals (average consistency of 52.1%; empirical *p* < 0.05; **Fig. 2**). Importantly, at more stringent *p*-value thresholds, as many as 91.4% of the “reciprocal” matched pairs had directional consistency, compared to a ceiling of 90.3% for the shared knowns, indicating that these matches for unknown signals show similar consistency as shared known metabolites. In addition, we estimated true positive rate for the matched signals by assuming that true matches would show same degree of consistency as the shared knowns, and found that the “multiple” setting generated more true positive matches overall, albeit at a lower confidence level (maximum true positive rate of 83.0%). In summary, the most stringent application of our method (“reciprocal” setting) identified ~7 times as many high-confidence matches compared to the shared knowns. The method can also be used to generate many more potential matches under more lenient settings (e.g. cross-method matching, **Supplementary Fig. 5**).

**Figure 2.**
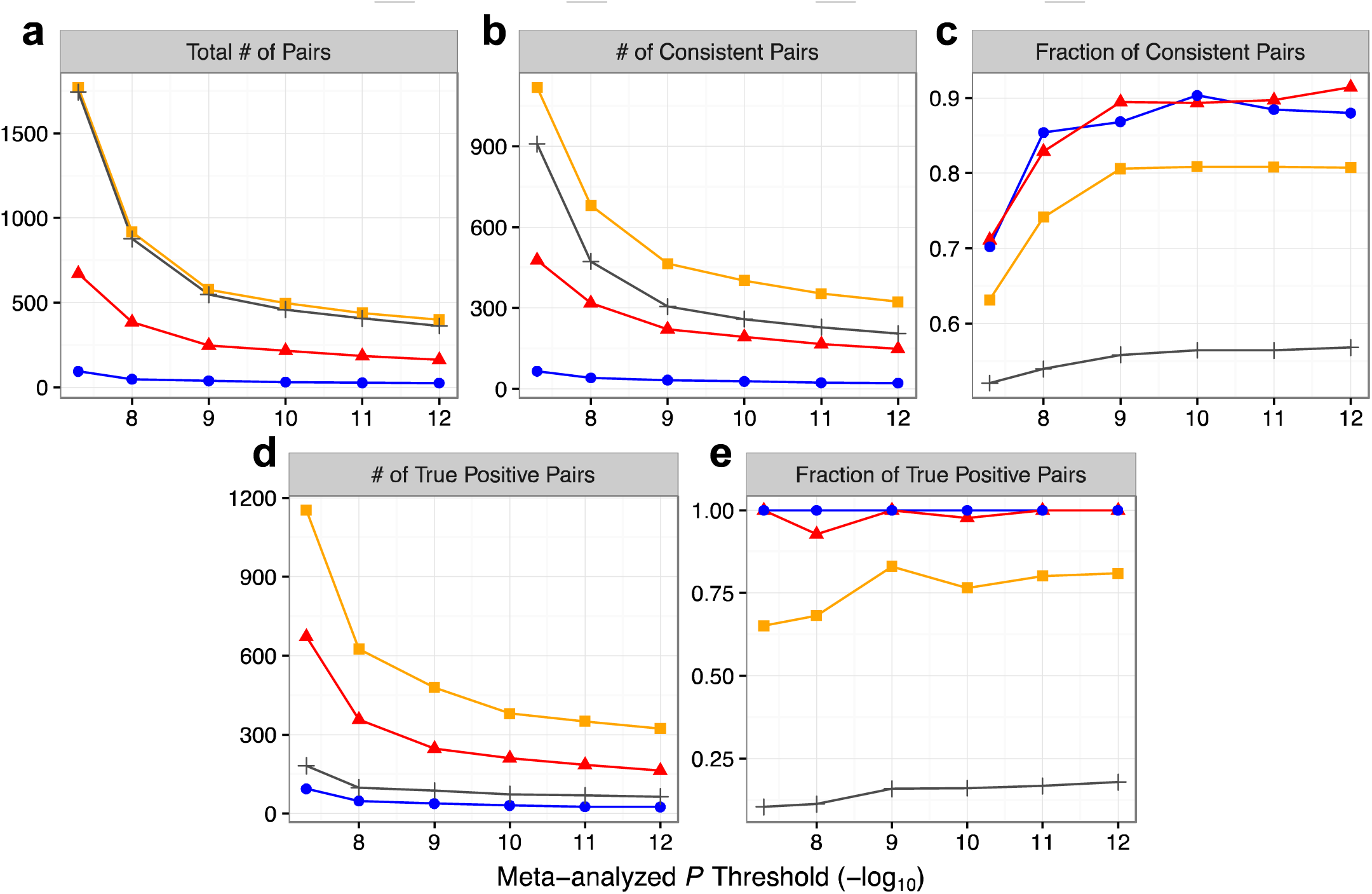
Genetic validation of OE-MCDS matched signals. GWAS were performed for “multiple” matched signal pairs (a subset of were “reciprocal”), shared known pairs (“Shared Known”, positive control), and randomly matched pairs (average statistics shown as “Random”, negative control) in OE and MCDS, followed by meta-analysis that ignored direction of effect. For each pair, the SNP with the best meta-analyzed *p*-value was selected to assess directional consistency of its association in the two cohorts. (a) “Total # of Pairs”: number of signal pairs with best SNPs below *p*-value threshold; (b) “# of Consistent Pairs” and (c) “Fraction of Consistent Pairs”: number and fraction of pairs with directionally consistent best SNPs below *p*-value threshold; (d) “# of True Positive Pairs” and (e) “Fraction of True Positive Pairs”: number and fraction of true positive pairs were estimated as described in Methods. X-axes start at genome-wide significant *p*-value threshold (*p* < 5 × 10^−8^). Error bars for “Random” pairs were excluded due to low visibility (all close to average).

#### Application to perform GWAS replication and meta-analysis

In order to directly assess how much power we gained from matching metabolite signals, we also performed GWAS replication analysis using OE and MCDS as the discovery and replication cohorts, respectively. Out of 14 shared knowns that had genome-wide significant associations in OE, 7 had nominally significant (*p* < 0.05) and directionally consistent associations in MCDS. In comparison, 156 “reciprocal” matched signals had genome-wide significant hits in OE, 56 of which had replicated associations in MCDS. Furthermore, after meta-analyzing the two cohorts (this time taking direction of effect into account), we found 71 shared knowns and 512 matched signals with genome-wide significant associations. Therefore, we identified ~7-8 times more metabolite signals with significant associations by using our method to pair up the unshared signals.

#### Application to identify BMI-associated signals

To further showcase the utility of signal matching, we applied our method to identify metabolite signals associated with BMI in OE and MCDS. First, we identified 1,474 signals (114 known and 1,360 unknown) that were significantly (*p* < 0.05/13,613) associated with BMI in OE and tested for their replication in MCDS. Out of the 96 BMI-associated OE known metabolites that were also measured in MCDS, 75 (78.1%) showed directionally consistent association with BMI at a nominal significance threshold in MCDS (**Supplementary Table 2**). For BMI-associated OE signals that could not be compared directly in MCDS (i.e. unknowns or unshared knowns), we applied our matching procedure, and found 322 out of 425 (89.2%) “reciprocal” or 543 out of 790 (68.7%) “multiple” matched signals to have directionally consistent and nominally significant association in MCDS. Thus, by matching signals across OE and MCDS, we replicated ~4-7 times more BMI-associated signals. Even if the unknown signals are not completely independent (i.e. multiple signals might correspond to one functional metabolite), the additional biological information extracted from the unknowns can be useful in pathway-based analyses that account for redundancy among signals, which we explored in the following section.

### Generating pathway annotations for metabolite signals

#### Pathway method overview and calibration

Metabolic pathway annotations in public databases contain rich information for characterizing metabolites. However, these annotations are usually binary (i.e. a metabolite is either in or out of a pathway with no uncertainty) and only involve metabolites with known identities. Thus, in order to predict functions of unknown signals in our data and to perform well-powered pathway analysis, we developed an approach to assign metabolite signals to previously curated pathways (**Fig. 1b**). First, we used profiling data from the BioAge cohort to construct “metabolic components” (MCs) that represent modules of covarying signals (**Supplementary Fig. 6**). Next, we collected metabolic pathway annotations from ConsensusPathDB (CPDB)^16^ and consolidated them into metabolite sets with unique metabolite combinations (i.e. one metabolite set may correspond to multiple pathways that contain identical sets of metabolites). We used the MCs to extend (reconstitute) the metabolite sets to include both known and unknown signals, resulting in a numeric signal-metabolite set annotation matrix, in which each signal is assigned a membership score in each set based on its similarity (across MCs) to metabolites originally curated to be in the set.

After reconstitution, we calculated a “label confidence score” to indicate how similar each reconstituted metabolite set is to the original pathways labeling the set; we also tested how well the reconstituted membership scores could be used to classify known metabolites into their original metabolite sets (see **Methods**). Using these performance measures, we calibrated different parameters in the reconstitution procedure (i.e. which metabolite sets and MCs to include) and picked the best-performing settings to construct the final BioAge annotation matrix (**Supplementary Fig. 7** and **Supplementary Table 3**).

#### Genetic validation of pathway annotations

We utilized OE genetic data to validate that the BioAge annotation matrix captures biologically relevant relationships. Specifically, we tested whether metabolite signals genetically associated with a gene would be enriched for reconstituted pathways related to that gene. First, we performed exome chip association analyses for all OE metabolite signals, mapped the signals with suggestive associations (*p* < 1 × 10^−5^) to BioAge using our matching procedure (in “multiple” setting), and derived a total of 6,465 associations between OE exome SNPs and BioAge matched signals. Next, we filtered the association results to select 36 loci that (1) contain coding SNPs associated with >= 5 signals and (2) contain genes curated to be in pathways reconstituted in the BioAge annotation matrix.

For each locus, we first used the BioAge annotation matrix to prioritize reconstituted metabolite sets enriched for the SNP-associated signals, and then checked whether the gene-related pathways showed up as enriched in the analysis. As negative control, we also repeated this procedure using 20 iterations of null association results. At an enrichment significance cutoff of *p* < 0.01, 5 out of 36 loci had gene-related enriched pathways (**Table 2**), compared to a mean of only 0.85 loci in null results (empirical *p* < 0.05; **Supplementary Table 4**). Even at less stringent enrichment significance thresholds, our approach performed better compared to null prioritization.

**Table 2.**
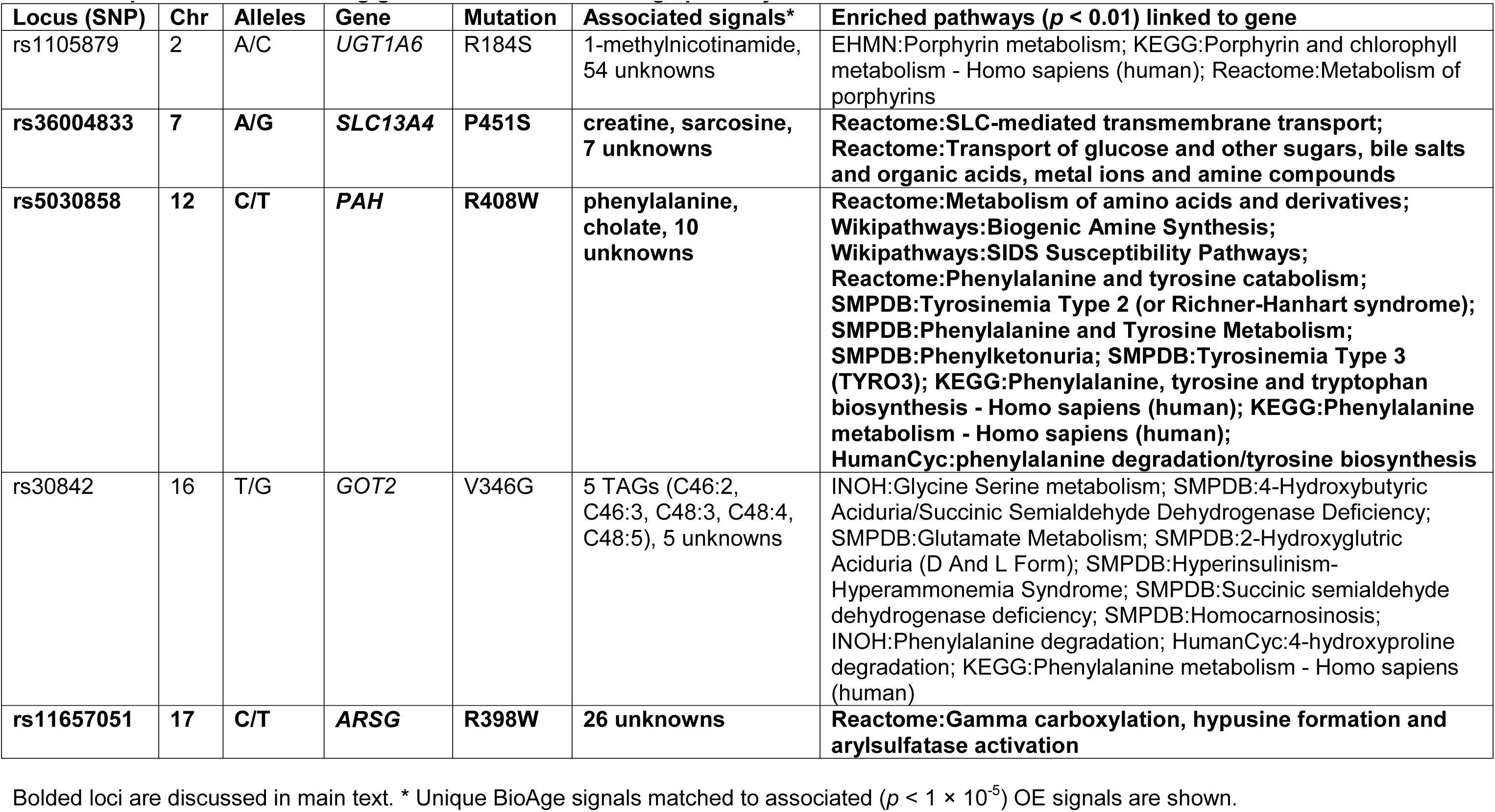
CTop loci identified during genetic validation of BioAge pathway annotations.

| **Locus (SNP)** | **Chr** | **Alleles** | **Gene** | **Mutation** | **Associated signals\*** | **Enriched pathways ($p < 0.01$) linked to gene** |
| --- | --- | --- | --- | --- | --- | --- |
| rs1105879 | 2 | A/C | *UGT1A6* | R184S | 1-methylnicotinamide, 54 unknowns | EHMN:Porphyryn metabolism; KEGG:Porphyryn and chlorophyll metabolism - Homo sapiens (human); Reactome:Metabolism of porphyrins |
| **rs36004833** | **7** | **A/G** | ***SLC13A4*** | **P451S** | **creatine, sarcosine, 7 unknowns** | **Reactome:SLC-mediated transmembrane transport; Reactome:Transport of glucose and other sugars, bile salts and organic acids, metal ions and amine compounds** |
| rs5030858 | 12 | C/T | *PAH* | R408W | phenylalanine, cholate, 10 unknowns | Reactome:Metabolism of amino acids and derivatives; Wikipathways:Biogenic Amine Synthesis; Wikipathways:SIDS Susceptibility Pathways; Reactome:Phenylalanine and tyrosine catabolism; SMPDB:Tyrosinemia Type 2 (or Richner-Hanhart syndrome); SMPDB:Phenylalanine and Tyrosine Metabolism; SMPDB:Phenylketonuria; SMPDB:Tyrosinemia Type 3 (TYRO3); KEGG:Phenylalanine, tyrosine and tryptophan biosynthesis - Homo sapiens (human); KEGG:Phenylalanine metabolism - Homo sapiens (human); HumanCyc:phenylalanine degradation/tyrosine biosynthesis |
| rs30842 | 16 | T/G | *GOT2* | V346G | 5 TAGs (C46:2, C46:3, C48:3, C48:4, C48:5), 5 unknowns | INOH:Glycine Serine metabolism; SMPDB:4-Hydroxybutyric Aciduria/Succinic Semialdehyde Dehydrogenase Deficiency; SMPDB:Glutamate Metabolism; SMPDB:2-Hydroxyglutric Aciduria (D And L Form); SMPDB:Hyperinsulinism-Hyperammonemia Syndrome; SMPDB:Succinic semialdehyde dehydrogenase deficiency; SMPDB:Homocarnosinosis; INOH:Phenylalanine degradation; HumanCyc:4-hydroxyproline degradation; KEGG:Phenylalanine metabolism - Homo sapiens (human) |
| **rs11657051** | **17** | **C/T** | ***ARSG*** | **R398W** | **26 unknowns** | **Reactome:Gamma carboxylation, hypusine formation and arylsulfatase activation** |

Bolded loci are discussed in main text. \* Unique BioAge signals matched to associated ( $p < 1 \times 10^{-5}$ ) OE signals are shown.

#### Examples of successful validation

We highlight three loci in which the enriched pathways have clear relevance to the affected genes’ functions. First, SNP rs5030858 causes a missense mutation (R408W) in *PAH*, which encodes the phenylalanine hydroxylase responsible for converting phenylalanine to tyrosine, and the mutation is known to cause phenylketonuria^17^. In our analysis, the SNP was associated with phenylalanine, cholate, and 10 unknown signals, which were enriched in 83 metabolite sets (corresponding to 129 pathways) at *p* < 0.01. Eight of these sets contain pathways directly linked to *PAH* in CPDB, while many others are also related to phenylalanine and tyrosine metabolism, such as those involved in amine, neurotransmitter, and tryptophan metabolism (**Table 2** and **Supplementary Table 5**).

Next, SNP rs36004833 is a missense variant (P451S) in *SLC13A4* and was associated with creatine, sarcosine, and 7 unknowns. *SLC13A4* encodes a sodium/sulfate cotransporter that is known to be expressed in the high endothelial venules^18^. Two of the metabolite sets enriched for the SNP-associated signals contain gene-related pathways (“Reactome: SLC-mediated transmembrane transport” and “Reactome: Transport of glucose and other sugars, bile salts and organic acids, metal ions and amine compounds”) (**Table 2** and **Supplementary Table 6)**. Importantly, creatine and sarcosine were not originally curated to be in these pathways, indicating that the pathways could not have been detected without using our method.

Finally, SNP rs11657051 causes a missense mutation (R398W) in *ARSG* (Arylsulfatase G^19^) and was associated with 26 unknown signals. The enriched metabolite sets contain one pathway directly linked to *ARSG* (“Reactome:Gamma carboxylation, hypusine formation and arylsulfatase activation”), demonstrating that our approach can be successful even without looking at any known metabolite associations (**Table 2** and **Supplementary Table 7**).

#### Application to identify BMI-associated pathways

As a last demonstration of the utility of PAIRUP-MS, we applied both the signal matching and pathway annotation methods to identify metabolic pathways associated with BMI. First, we identified 1,474 and 1,289 signals associated with BMI in OE and MCDS, respectively (after correction for multiple testing), and matched the unshared signals from each cohort to a common reference cohort (BioAge) using the “multiple” match type setting. Between shared knowns and matched signals, we could map a total of 1,162 OE and 1,030 MCDS BMI-associated signals to BioAge. We performed separate pathway enrichment analyses using these two sets of signals and the BioAge reconstituted metabolite sets. In total, 218 (31.6%) and 179 (25.9%) out of 690 metabolite sets were enriched (FDR <= 5%) for the OE-and MCDS-matched signals, respectively (**Supplementary Table 8**). A total of 107 (15.5%) metabolite sets, corresponding to 215 pathways, were enriched in both analyses (overlap significance: Fisher’s *p* = 4.01 × 10^−20^; **Fig. 3**). These pathways are involved in a wide range of biological processes, including signal transduction, neurotransmission, synaptic function, macronutrient metabolism, immune system, and transport of various compounds.

**Figure 3.**
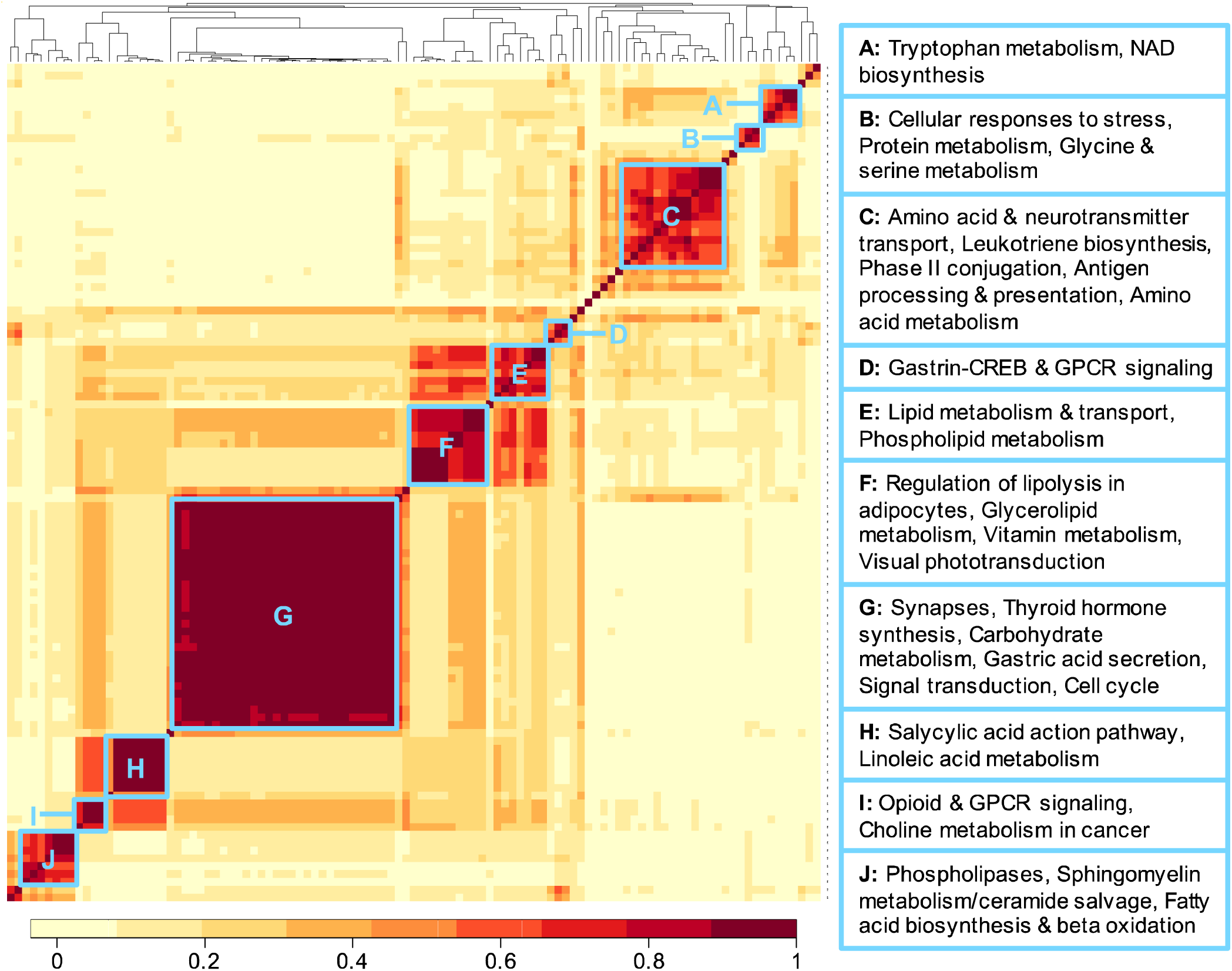
Correlation heat map of 107 metabolite sets enriched (5% FDR) for both OE and MCDS BMI-associated signals. Correlation between metabolite sets were calculated using the BioAge annotation matrix. Color key indicates correlation values between each pair of metabolite sets. Distinct clusters (blue boxes) are labeled with representative pathway names in the clustered metabolite sets.

To show that our methods increased power for detecting BMI-related pathways, we performed traditional overrepresentation analyses using only the BMI-associated known metabolites and the binary metabolite set annotations from CPDB, and found only 25 overrepresented sets for OE at 5% FDR, while MCDS had no significant sets at all (**Supplementary Table 9**). Overall, these results showcase how PAIRUP-MS could be used to incorporate unknown signals into pathway-based analyses, improving both power and interpretation in metabolomics studies.

## DISCUSSION

While untargeted profiling generate data for thousands of metabolite signals, most signals cannot be readily identified as known metabolites, are difficult to compare across studies, and consequently are often excluded from downstream analyses. In order to facilitate better use of untargeted data, we have developed PAIRUP-MS, an integrative tool for processing, matching, and annotating the unknown signals, which allowed us to compare them across MS-based datasets, extract useful information from their data, and improve power at detecting trait-associated biology. To our knowledge, no existing tools provide a similar streamlined and flexible framework for dealing with unknown signals across multiple datasets.

Within PAIRUP-MS, we implemented a m/z and imputation-based method for matching unknown signals across datasets, demonstrated that it could be used to accurately pair up shared known metabolites, and showed that it could outperform a m/z and RT-based matching approach when RT information is unreliable. When we used genetic associations to validate a subset of the matched signals, we observed that the most stringent matches performed comparably in validation to the shared knowns. By including matched signals in combined association analyses (with genetic variants or with phenotypes), we were able to identify ~7 times more signals with significant replicated associations compared to the common practice of only studying known metabolites.

We also developed a pathway reconstitution procedure to annotate unknown signals using curated metabolic pathways. Again, we used genetic data to validate the pathway annotations, showing that signals associated with a gene were likely to be enriched for the gene-related pathways. Finally, we applied both the matching and pathway methods to identify BMI-associated pathways using a crossover approach (i.e. identifying trait-associated signals in one dataset but testing for enrichment using pathways built from another dataset), and demonstrated that it was more powerful than performing pathway overrepresentation analysis using only known metabolites.

Several limitations of this study should be considered. First, although the samples encompassed a range of ascertainments and were collected in different continents, all three datasets were generated by the same LC-MS profiling platform (albeit over different iterations of that platform). Thus, it will be valuable to test PAIRUP-MS using data from other platforms. The pathway method should be applicable to any profiling dataset that has sufficient number of identified known metabolites, while the matching method is applicable to any MS-based dataset. However, the performance of the methods, especially the matching of unknown signals, may be influenced by technical differences across platforms, so further validation will be important. For both methods, calibrating parameters using known metabolites, as we have described here, will be crucial to achieving good results when analyzing data from diverse studies and platforms.

One consequence of using known metabolites for calibration is that analyses may be more robust for metabolite types that are well represented in the known metabolites or are well-annotated in pathways. This problem is more severe for the pathway method, in which we also filtered and reconstituted the metabolite sets using the knowns. Hence, even though the main goal of our methods is to analyze the unknown signals, progress in known metabolite identification and pathway curation can greatly improve the performance of our methods. Conversely, our approach may also be applied to facilitate the process of metabolite identification and expand knowledge on previously curated pathways.

Another limitation of our study is that validation of both methods required genetic data to be available for the profiled samples. In particular, we were only able to validate OE-MCDS matched signals because BioAge genotypes were unavailable, and even the OE and MCDS cohorts had ancestry differences that complicated the interpretation of results (i.e. hard to determine if failed validation was due to false match or genetic difference). Larger numbers of untargeted profiling datasets with accompanying genetic data will allow the ongoing use of genetic data to assess the performance of our methods across a wider range of samples.

There are several major directions for applying and extending our methods in the future. First, while we focused on analyzing pairs of datasets in this paper, large-scale meta-analysis requires a framework to match and compare many datasets simultaneously. Our current recommendation would be to define a “reference panel” dataset and match metabolite signals in all other datasets to this common reference (analogous to reference panels for genotype imputation). The choice of the reference panel could be made after assessing the performance of matching shared known metabolites across pairs of datasets. A reference dataset that incorporates diverse samples and captures a wide range of metabolite abundance variation would also be ideal for building reference pathway annotations that can be used to annotate all signals that are matched to the reference.

Furthermore, we note that we focused exclusively on metabolite profiling data from plasma samples. It would be interesting to apply PAIRUP-MS to datasets extracted from other types of biological samples. In particular, it would be of interest to test how the pathway reconstitution method could be used across different tissues to capture tissue-specific patterns of metabolic variations, analogous to the value of tissue and cell type-specific gene expression measurements for gene set reconstitution^20^.

In conclusion, we have developed a suite of methods, PAIRUP-MS, for analyzing unknown signals in untargeted profiling data and showed that it can be applied to improve power and make biological inferences in metabolomics analyses. Matching across datasets is vital for increasing power through meta-analysis (e.g. imputation enabled meta-analysis of GWAS data across different genotyping platforms). Similarly, PAIRUP-MS can enable much more powerful analyses and meta-analyses of the great majority of untargeted metabolomics data that has not yet been systematically examined. The availability of many more untargeted datasets (especially with accompanying genetic data) will allow these analyses to be performed in much larger sample sizes, and should also allow further refinement and evaluation of our methods across a wider range of datasets, samples, and platforms.

## METHODS

### Metabolomics datasets and data processing

#### Obesity Extremes cohort (OE)

Metabolite profiling was performed on plasma samples of 300 individuals from the Estonian Biobank of the Estonian Genome Center at the University of Tartu (EGCUT)^21^. The individuals were selected from the extremes of the BMI distribution to include 100 lean (BMI < 20), 100 obese (BMI > 34), and 100 random individuals matched for age, sex, and fasting time (>= 4 hr). Clinical data were examined to exclude any individuals with pregnancy, anorexia, or known wasting illnesses. Profiling was done using 4 LC-MS methods to measure 393 known metabolites and 18,667 unknown signals.

#### Mexico City Diabetes Study cohort (MCDS)

Metabolite profiling was performed on fasting plasma samples collected from 865 individuals in the Mexico City Diabetes Study at the 1998 exam cycle. Details of this prospective cohort have been described previously^22^. 3 LC-MS methods were used to measure 306 known and 11,888 unknown signals.

#### BioAge Labs Mortality cohort (BioAge)

Metabolite profiling was performed on plasma samples of 583 individuals 70-80 years old and free of major morbidity at enrollment, selected from the Estonian Biobank as a mortality cohort. None of the BioAge samples overlap with OE. 4 LC-MS methods were used to measure 629 known and 19,087 unknown signals.

#### Metabolite data processing

We performed the following steps on each of the 3 metabolite datasets before using them in downstream analyses: (1) log-transformed and normalized the data using internal standards and interspersed pooled plasma samples, (2) removed outlier data points and signals with noisy variation trends, (3) removed samples and signals with > 25% (OE) or 50% (MCDS and BioAge) missing data, (4) imputed missing values using the MICE R package (v2.25)^23^, (5) adjusted each signal for covariates if appropriate (only done for data used in genetic and BMI association analyses: age, sex, and fasting time for OE; age and sex for MCDS), and (6) performed rank-based inverse normal transformation to calculate abundance *z*-scores for each signal.

More details of the metabolite profiling, quality control, and missing value imputation procedures are described in **Supplementary Information**.

### Matching metabolite signals across datasets

#### Checking for m/z agreement

We implemented an algorithm to check whether the difference between 2 m/z values are within *d* ± 0.005 of each other. While *d* = 0 in the most basic setting, it can be set to multiple values when taking ionization modes and adduct ions into account as shown in **Supplementary Table 10**. In particular, we implemented 2 settings when at least one of the m/z values was derived from positive ion mode: (1) “No adduct”: only the most common positive adduct ion is considered, or (2) “Adduct”: 4 common positive ions (M^+^, [M+H]^+^, [M+NH_4_]^+^, and [M+Na]^+^) are considered. When identifying all signal pairs with agreeing m/z values across two datasets, we also implemented a third “combined” setting, where the “adduct” setting was used only when the “no adduct” setting failed to identify any pairs with matching m/z values. We tested all 3 settings in *Parameter calibration* to identify the optimal setting to use for matching signals across each pair of datasets.

#### Calculating imputation-based correlation

To impute a signal from Dataset 1 to Dataset 2, we fit a linear regression model for the signal using Dataset 1 data, which included all known metabolites shared by the two datasets as predictors, and then applied the resulting model to Dataset 2 data to predict the signal’s abundance in Dataset 2 samples. The reversed process was done to impute signals from Dataset 2 to Dataset 1. When performing imputation for the shared knowns (used as positive controls in *Parameter calibration*), we built a leave-one-out model for each known metabolite *m*, where all shared knowns except for *m* were used as predictors. After imputation, we calculated pairwise correlation between Dataset 1 and Dataset 2 signals using 3 different settings: (1) “Dataset 1 correlation”: correlate Dataset 1 (observed) and Dataset 2 (imputed) signals across Dataset 1 samples, (2) “Dataset 2 correlation”: correlate Dataset 1 (imputed) and Dataset 2 (observed) signals across Dataset 2 samples, or (3) “All correlation”: merge observed and imputed data and correlate signals across all samples. We evaluated these settings in *Parameter calibration* to identify the optimal approach to use for *Matching based on m/z and correlation*.

#### Matching based on m/z and correlation

In general, to match a Dataset 1 signal, *s*, to a Dataset 2 signal, we would first identify all Dataset 2 signals whose m/z agree with *s*, and then select the one that is best correlated with *s* to be the final match. If no Dataset 2 signals share matching m/z with *s*, then *s* is not matched at all. We implemented variations of this procedure and compared their performance in *Parameter calibration*. First, since the matching procedure is directional (i.e. matching from Dataset 1 to 2 vs. 2 to 1 could be different), we distinguished between 3 types of matches, each type being a subset of the previous type: (1) “Multiple”: all matches that can be made as described by the general guideline above, which allows multiple-to-1 matching (i.e. multiple Dataset 1 signals can be mapped to the same Dataset 2 signal), (2) “Unique”: unique 1-to-1 matches that only keep the best (in terms of correlation) Dataset 1 signal mapped to each Dataset 2 signal, or (3) “Reciprocal”: unique, bidirectional 1-to-1 matches that appear both when matching from Dataset 1 to 2 and from 2 to 1. We also allowed 2 partition settings: (1) “Within method”: only look for matches among signals measured using the same profiling method, or (2) “Across method”: look for matches across datasets regardless of differences in profiling methods. Lastly, we implemented an optional correlation cutoff for calling matches (i.e. only keep matched pairs whose correlation is greater than the cutoff).

#### Matching based on m/z and RT

We implemented a crude RT-based matching procedure to serve as a comparison to our imputation-based method. Here, to match signals from Dataset 1 to Dataset 2, we would first use RTs of shared knowns to fit a linear model for predicting the RT shift between the two datasets. Then, to match a Dataset 1 signal, *s*, to a Dataset 2 signal, we would predict the RT of *s* in Dataset 2 using the fitted model, find all Dataset 2 signals whose m/z agree with *s*, and then select the one whose RT is closest to the predicted RT to be the final match. Just as we did for *Matching based on m/z and correlation*, we tested different match types (“multiple”, “unique”, or “reciprocal”) and partition settings (“within method” or “across method”) in *Parameter calibration*.

#### Parameter calibration

To calibrate all of the parameters described above (i.e. adduct ion setting, correlation setting, match type, partition setting, and correlation cutoff), we treated known metabolites shared across two datasets as unmatched signals and applied our procedures to match them up. For each pair of datasets (i.e. OE-MCDS, OE-BioAge, or MCDS-BioAge), we calculated the number and percentage of (1) correct matches: Dataset 1 metabolite is matched to itself in Dataset 2, (2) highly correlated matches: Dataset 1 metabolite is matched to a signal that is highly correlated (r^2^ > 0.8) with itself in Dataset 2, and (3) incorrect but highly correlated matches: matches in (2) but not in (1).

### GWAS and genetic validation of matched signals

#### GWAS and meta-analysis

294 OE samples with Estonian reference imputed genotypes and 637 MCDS samples with 1000 Genomes phase 3 imputed genotypes were used to perform GWAS for the OE-MCDS shared known and matched signal pairs (see **Supplementary Information** for genotyping and imputation details). We used the linear mixed model and kinship matrix calculation methods in EPACTS (v3.2.6)^24^ to analyze each cohort, restricting analyses to biallelic SNPs with minor allele count >= 3 in the cohort and including genotyping platform as a covariate. SNPs that overlap between the 2 cohorts were meta-analyzed using the inverse variance weighted method in METAL (2011-03-25 version)^25^.

#### Validation analysis

For each pair of shared knowns or matched signals, we performed an alternative meta-analysis that ignored direction of effect (i.e. forcing all effect size estimates to be positive), identified the SNP with the best meta-analyzed *p*-value, and determined if its original effect size estimates in OE and MCDS were directionally consistent. We combined results for individual signal pairs to calculate the number and fraction of consistent pairs among shared knowns or matched signals at different *p*-value thresholds (i.e. only considering pairs whose best SNPs pass the threshold). We used Fisher’s exact test to evaluate statistical significance of the observed degree of consistency. Additionally, for the matched signals, we performed 20 null permutations by shuffling the mapping between matches (i.e. generating randomly matched pairs), and then calculated an empirical *p*-value for the observed consistency fraction (i.e. *p* = proportion of null fractions greater than or equal to the observed fraction). Finally, we estimated the fraction of true positive pairs (i.e. true positive rate) at each *p*-value threshold to be: max[(*c* – 0.5)/(*s* – 0.5), 1] where *c* = fraction of consistent pairs and *s* = fraction of consistent shared knowns, and then derived the number of true positive pairs using this fraction.

### Generating pathway annotations for metabolite signals

#### Metabolic pathway and metabolite set annotations

We collected metabolic pathway annotations from CPDB (release 31)^16^, which aggregated data from 11 source databases. We filtered the annotations to obtain pathways containing >= 2, 5, or 10 known metabolites profiled in BioAge and assigned a “*database*:*name*” label to each pathway, where *database* refers to the source database and *name* is the pathway name as shown in that database. We further consolidated the pathways into metabolite sets, with each set representing all pathways sharing the same metabolite combination, excluding metabolites not profiled in BioAge. The resulting metabolite set annotations were used as input data in *Signal-metabolite set annotation matrix*. CPDB also contains annotations that link genes to pathways based on previous literature evidence, which we utilized in *Genetic validation of pathway annotations*.

#### Metabolic components

We calculated pairwise Spearman correlation for all metabolite signals (known and unknown) across BioAge samples, and then performed principal component analysis on the correlation matrix to derive “metabolic components” (MCs; eigenvectors that represent modules of correlated signals) and a corresponding signal-MC score matrix. In order to assess if the MCs captured biological patterns contained within metabolite sets derived from CPDB, we performed enrichment analysis using the MC scores of known metabolites included in the metabolite sets. For each MC and each set, we performed a two-tailed Wilcoxon rank-sum test to compare the MC scores of metabolites in the set vs. other metabolites. We repeated the tests using 20 iterations of null, permuted MC scores and estimated the FDR for each observed rank-sum *p*-value threshold, *P*, to be: (average number of null *p*-values <= *P* per iteration) / (number of observed *p*-values <= *P*). This FDR was used to filter MCs as described in *Parameter calibration*.

#### Signal-metabolite set annotation matrix

We used the MC scores of metabolite signals to reconstitute the CPDB metabolite sets, adopting a previously described framework for gene set reconstitution^20^. Again, we performed a two-tailed Wilcoxon rank-sum test to compare the MC scores of known metabolites in the set vs. not in the set for each metabolite set-MC pair, but this time storing the results as a metabolite set-MC matrix of rank-sum *z*-scores. Next, we used the signal-MC matrix (from *Metabolic components*) and the metabolite set-MC matrix to calculate the Spearman correlation between each signal-metabolite set pair across the MCs, resulting in a signal-metabolite set annotation matrix of correlation scores. To avoid overfitting, when calculating the matrix scores of a known metabolite originally annotated in a set, we repeated the reconstitution procedure by leaving out the data for this metabolite.

#### Parameter calibration

A calibration procedure was carried out to determine which metabolite sets and MCs to use for constructing the optimal signal annotation matrix. We tested metabolite sets containing >= 2, 5, or 10 known metabolites profiled in BioAge. For MCs, we tested the top 50, 100, or 583 MCs (in terms of variance explained), or filtered for any MCs that were enriched for >= 1 metabolite set at 1% or 5% FDR (see *Metabolic components*). After generating annotation matrices using different parameters, for each matrix, we converted correlation scores of known metabolites into *p*-values, and then evaluated how well these *p*-values could be used to classify metabolites into their original metabolite sets using a receiver operating characteristic (ROC) curve and area under the curve (AUC) approach. We also performed post-reconstitution enrichment analysis to identify reconstituted metabolite sets enriched for metabolites originally annotated to be in the set (see *Metabolite set enrichment analysis*). The enrichment *p*-values from this analysis were used as “label confidence scores” for the metabolite sets.

### Genetic validation of pathway annotations

#### Exome chip association analysis

A subset of 281 OE samples were genotyped using the Illumina HumanOmniExpressExome BeadChip (v1.2). We performed association analysis for each OE metabolite signal using this genetic dataset to identify exome SNPs associated with signal abundance. We restricted our analyses to 57,220 exome SNPs with minor allele count >= 3 and call rate >= 0.5 in OE, using the linear mixed model and kinship matrix calculation methods in EPACTS (v3.2.6)^24^.

#### Validation analysis

We filtered the OE exome chip association results using OE-BioAge “multiple” matched signals to identify associations between OE SNPs and BioAge signals. We grouped the associations by SNPs and identified SNPs that (1) were associated with >= 5 signals at *p* < 1 × 10^−5^, (2) cause missense or nonsense mutations in genes, and (3) affect genes linked to >= 1 CPDB pathways used for reconstitution. If multiple SNPs were found for the same gene, we only kept the SNP with the best *p*-value. To perform validation analysis for each locus, we first used the BioAge signal annotation matrix to prioritize metabolite sets enriched for the SNP-associated signals (see *Metabolite set enrichment analysis*). Next, we determined if any pathways linked to the affected gene showed up as enriched across a range of enrichment *p*-value thresholds. For each threshold, we counted the number of loci for which >= 1 gene-associated pathway was enriched, and then repeated the procedure using 20 iterations of null exome chip association results (i.e. identifying top signals associated with random genotypes) to generate null counts for comparison. We calculated the empirical *p*-value for the observed count to be the proportion of null counts >= the observed count.

### BMI-associated metabolite signals and metabolite sets

#### Metabolite signals

We calculated BMI *z*-scores for both OE and MCDS samples by adjusting raw BMI for age and sex and performing rank-based inverse normal transformation on the residuals. For OE, all samples in the Estonian Biobank were included during *z*-score calculation; for MCDS, only samples in the cohort were included. We performed linear regression to test for association between BMI *z*-scores and signal abundance *z*-scores in both cohorts and identified BMI-associated signals at Bonferroni-corrected significance thresholds (*p* < 0.05/13,613 for OE; *p* < 0.05/7,136 for MCDS).

#### Metabolite sets

We identified BioAge shared knowns or “multiple” matched signals corresponding to the BMI-associated OE and MCDS signals from the previous section, and then used these signals and the BioAge signal annotation matrix to perform two separate enrichment analyses as described in *Metabolite set enrichment analysis*. After identifying enriched metabolite sets at 5% FDR in both analyses, we performed a Fisher’s exact test to evaluate the statistical significance of the overlap.

### Metabolite set enrichment analysis

#### Rank-sum p-value

To calculate the enrichment of each metabolite set in the signal annotation matrix given a “positive” and a “negative” list of signals, we used a two-tailed Wilcoxon rank-sum test to compare the metabolite set scores (absolute correlation) of the positive vs. negative signals to get a nominal *p*-value. For post-reconstitution enrichment analysis (in *Generating pathway annotations for metabolite signals*), the positive and negative lists consisted of known metabolites in an original CPDB metabolite set vs. not in the set, respectively. For identifying trait-associated metabolite sets, the lists consisted of signals significantly associated with the trait (SNP genotype in *Genetic validation of pathway annotations*; BMI in *BMI-associated metabolite signals and metabolite sets*) vs. all other signals (restricting to signals that could be matched across datasets).

#### Permutation p-value

For *Genetic validation of pathway annotations* and *BMI-associated metabolite signals and metabolite sets*, we generated null lists of signals ranked by their association with 1000 sets of permuted trait scores and repeated the rank-sum tests using the null lists (i.e. taking the top *N* signals in each null list to be the “positive” list, where *N* = the number of signals in the observed positive list). We used the null enrichment results to adjust the observed rank-sum *p*-values for metabolite set-specific biases, calculating a permutation *p*-value for each metabolite set to be the proportion of null rank-sum *p*-values <= the observed rank-sum *p*-value.

#### FDR

For *BMI-associated metabolite signals and metabolite sets*, we implemented another level of permutations to estimate analysis-wide FDR that accounts for multiple hypothesis testing across metabolite sets. We generated 20 additional null signal lists and calculated their corresponding rank-sum *p*-values. By comparing these new null rank-sum *p*-values against the 1000 sets of null rank-sum *p*-values computed previously, we calculated 20 sets of null permutation *p*-values. The FDR for an observed permutation *p*-value threshold, *P*, was then estimated as: (average number of null permutation *p*-values <= *P* per null set) / (number of observed permutation *p*-values <= *P*). Finally, we forced the final FDR estimates to be monotonic (i.e. FDR for smaller *p*-values had to be <= FDR for larger *p*-values) to smooth out random fluctuations in null iterations.

A schematic overview of how we calculated statistical significance in metabolite set enrichment analysis is shown in **Supplementary Figure 8**.

### Metabolite set overrepresentation analysis

Binary metabolite set annotations from CPDB were used to perform overrepresentation analysis given a list of trait-associated known metabolites. For each metabolite set, we created a 2 × 2 contingency table that categorized known metabolites based on trait association and metabolite set membership and used it to calculate a one-sided Fisher’s exact *p*-value. Null permutations were performed as described in *Metabolite set enrichment analysis* (except using Fisher’s exact test instead of rank-sum test) to calculate permutation *p*-value and FDR.

### Code availability

PAIRUP-MS source code and documentation are available at https://github.com/yuhanhsu/PAIRUP-MS.

## ACKNOWLEDGMENTS

We thank C. Astley, C. Clish, A. Ganna, A. Kamburov, R. Salem, and S. Vedantam for discussions on study design and/or methodology. We thank the Broad Metabolomics Platform and SIGMA T2D Consortium for sharing data resources. This work was supported by grants from the National Heart, Lung, and Blood Institute (F31HL126581), National Institute of Diabetes and Digestive and Kidney Diseases (R01DK075787), Doris Duke Charitable Foundation (215205), Estonian Research Council (IUT20-60, PUT1665, PUT1660), European Union through Horizon 2020 (692145), and European Union through the European Regional Development Fund (Project No. 2014-2020.4.01.15-0012).

## AUTHOR CONTRIBUTIONS

Y.H.H., C.C., T.H.P., T.E., and J.N.H. conceived and designed the methods; Y.H.H. and C.C. implemented the methods; J.M.M., A.M., K.Fisher, K.Fortney, E.K.M., C.G., M.E.G., and T.E. provided metabolite and/or genetic data for the study cohorts; Y.H.H. and T.E. performed the analyses; Y.H.H., T.H.P., and J.N.H. wrote the manuscript with feedback from all authors.

